# From CFTR to a CF signalling network: a systems biology approach to study Cystic Fibrosis

**DOI:** 10.1101/2023.11.15.567166

**Authors:** Matthieu Najm, Loredana Martignetti, Matthieu Cornet, Mairead Kelly-Aubert, Isabelle Sermet, Laurence Calzone, Véronique Stoven

**Affiliations:** Center for Computational Biology (CBIO), Mines Paris-PSL, Paris, 75006, France; Institut Curie, Université PSL, Paris, 75005, France; INSERM U900, Paris, France; Institut Necker Enfants Malades, INSERM U1151, Paris, 75005, France; Université Paris Cité, Paris, France; Centre de Référence Maladies Rares, Mucoviscidose et Maladies Apparentées, Hôpital Necker Enfants Malades AP-HP Centre Paris Cité, Paris, France

**Keywords:** Cystic Fibrosis, CF cellular phenotypes, CF signalling network, network topology, therapeutic target

## Abstract

**Background:** Cystic Fibrosis (CF) is a monogenic disease caused by mutations in the gene coding the Cystic Fibrosis Transmembrane Regulator (CFTR) protein, but its overall physio-pathology cannot be solely explained by the loss of the CFTR chloride channel function. Indeed, CFTR belongs to a yet not fully deciphered network of proteins participating in various signalling pathways.

**Methods:** We propose a systems biology approach to study how the absence of the CFTR protein at the membrane leads to perturbation of these pathways, resulting in a panel of deleterious CF cellular phenotypes.

**Results:** Based on publicly available transcriptomic datasets, we built and analyzed a CF network that recapitulates signalling dysregulations. The CF network topology and its resulting phenotype was found to be consistent with CF pathology.

**Conclusion:** Analysis of the network topology highlighted a few proteins that may initiate the propagation of dysregulations, those that trigger CF cellular phenotypes, and suggested several candidate therapeutic targets. Although our research is focused on CF, the global approach proposed in the present paper could also be followed to study other rare monogenic diseases.

## 1 Background

Cystic fibrosis (CF) is the most common life-limiting autosomal disease in the Caucasian population, affecting about 162.000 patients worldwide, of which 105.000 are diagnosed (Guo et al., 2022). It is caused by mutations in the *CFTR* gene encoding for the cystic fibrosis transmembrane conductance regulator (CFTR) protein, a chloride ion channel expressed at the apical membrane of polarized epithelial cells (Seibert et al., 1997). More than 2000 mutations in *CFTR* have been reported, but the deletion of the F508 amino-acid (F508del) is present in 70% of the mutated alleles in the Caucasian population, and most of the mutations lead to compromised transepithelial anion conductance (Veit et al., 2016). Various organs are affected in CF, but the most severe symptoms are in the lungs, where the defective chloride transport leads to the dehydration of surface mucus, chronic bacterial infection, and inflammation, causing lung tissue damage and ultimately, respiratory insufficiency.

However, CF symptoms not only result from the loss of CFTR-mediated anion conductance, but also from perturbations of other CFTR-dependent biological functions (Hanssens et al., 2021). Indeed, CFTR belongs to a protein-protein interactions (PPI) network (Pereira et al., 2021; Farinha and Gentzsch, 2021), and the absence of CFTR may perturb its direct or indirect interactors, and propagate dysregulations towards various biological pathways in which these interactors play a role. In agreement with this idea, studies on *CFTR* -/- knockout mice (Crites et al., 2015), *CFTR* -/- knockout piglets (Fleurot et al., 2022), and cell lines in which CFTR is inactivated by the CRISPR/Cas9 technology (Hao et al., 2020) have reported that the absence of CFTR affects cell signalling and transcriptional regulation. These dysregulations may explain various and apparently unrelated cellular phenotypes, including uncontrolled pro-inflammatory response (Jacquot et al., 2008), unbalanced oxidative stress with increased reactive oxygen species (Jeanson et al., 2012), impaired epithelial regeneration (Conese and Di Gioia, 2021), or perturbation of cell junctions and cytoskeleton (Pankonien et al., 2022).

To explain these seamlessly unrelated phenotypes, we propose to use a systems biology approach for CF, where the two aims are (1) to explore how the absence of CFTR can be functionally related to the signalling dysregulations that ultimately lead to CF cellular phenotypes; and (2) to suggest new therapeutic targets that may modulate these phenotypes.

Indeed, systems biology approaches provide tools for building network models to reason on complex systems. Subsequent topological analysis or dynamic mathematical models performed on these networks allow to study how different biological components of the networks interact to produce phenotypic properties, which is relevant to the questions at hand.

Systems biology approaches have seldom been implemented in monogenic diseases, but have been widely used in cancer, often referred to as a network disease (Hornberg et al., 2006), where intricate processes contribute to the emergence of unexpected and often non-intuitive phenotypes. Very few contributions have been devoted to systems biology approaches of CF. Previous studies have focused on the construction of the CFTR interactome that distinguishes PPI networks involving wt-CFTR and those involving the most frequent mutant F508del-CFTR (Pankow et al., 2015; Pereira et al., 2021). The latter led to the construction of a navigable knowledge map, the CyFi- MAP, that integrates all proteins known to be involved in the processing, maturation, retention and degradation of wt-CFTR and F508del-CFTR. Although the CyFi-MAP represents a key contribution for the problem of rescuing F508del-CFTR, this map does not tackle the questions of interest in the present paper. Other studies highlighted links between CFTR and signalling pathways involved in the disease (see (Pankonien et al., 2022) for a review), but they did not provide a global view of how CFTR is linked to dysregulated molecular mechanisms and to CF phenotypes. Recently, transcriptomic data have been produced to identify differentially expressed genes in CF. These genes were connected within a PPI network, based on information available in PPI databases (Trivedi et al., 2023). Although this network comprises genes that are consistent with current knowledge in CF, it does not contain CFTR, which prevents understanding the functional link between CFTR and the differentially expressed genes, or with CF cellular phenotypes.

To overcome the limitation of previous studies, in the systems biology approach proposed here, we build a comprehensive signalling network, called the CF network in the following, that recapitulates CF pathway dysregulations, using transcriptomic data available for CF and control patients and information available in biological pathway databases. As detailed below, we connected CFTR to this network based on PPI information. Analysis of the CF network topology allows to formulate hypotheses on key proteins and molecular mechanisms that functionally link CFTR to major CF cellular phenotypes, and to highlight potential targets that may counteract these phenotypes.

## 2 Results

### 2.1 Global approach to building the CF network

In systems biology, various networks can be built to represent different types of biological information, such as gene regulatory networks, genetic interaction networks, signal transduction networks, metabolic networks, PPI networks, or disease networks.

There is no universal technique that can be followed to construct networks, and the choice of their representation needs to be adapted to the question of interest. In the present study, we wish to establish a link between the absence of CFTR and the over- all signalling dysregulations leading to the cellular phenotypes that characterize CF. Therefore, we chose to build a CF network focusing on the signalling pathways that are perturbed in the disease, and where dysregulations in one pathway may affect other pathways. In order to avoid potential bias in the CF literature, we adopted a data-driven approach based on publicly available transcriptomic studies. We are aware that some CF phenotypes might arise from biological events that are not detectable in the transcriptome of CF cells, but we considered that gene expression data had the potential to capture some of the major molecular dysregulations present in CF cells. Our study relies on a meta-analysis of public transcriptomic datasets for CF respiratory epithelial cells and their Non-Cystic Fibrosis (NCF) control counterparts, allowing the identification of the signalling pathways dysregulated in CF. Based on information available in pathway databases, these dysregulated pathways share many common proteins, which allowed to connect them into a network. As detailed below, CFTR was absent from this network, because it did not belong to any of the differentially expressed signalling pathways. However, we observed that several proteins of the network were also present in the CFTR PPI interactome, either as direct interactors of CFTR, or as indirect interactors of CFTR via a single intermediate protein. This important result was consistent with the assumption that CFTR direct interactors may be perturbed in CF, and initiate the propagation of dysregulations within the CF network.

The figure 1 summarizes the global approach followed in the present study.

**Fig. 1.**
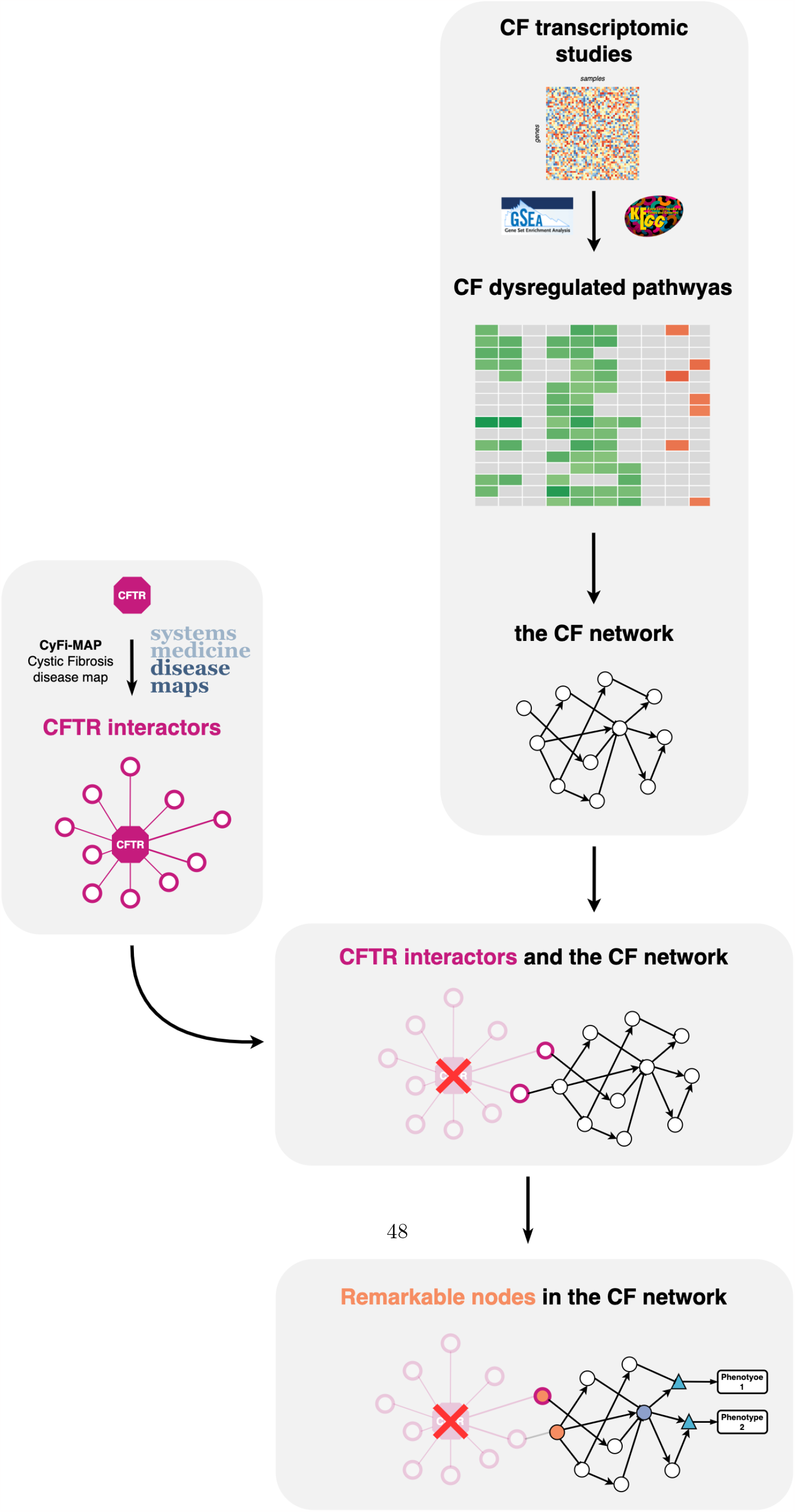
Global approach followed to build the CF network: A meta-analysis of CF transcriptomic data allowed the identification of dysregulated pathways and the construction of the corresponding CF network. This network comprises known CFTR interactors that can be viewed as source nodes initiating the propagation of dysregulations.

### 2.2 Selection of publicly available transcriptomics data

Many transcriptomic studies have been performed in CF over the last 15 years (Ideozu et al., 2019). However, these data suffer from a few limitations that are obstacles to improve our understanding of CF. First, they consider a wide range of cell types, including native nasal or bronchial cells, primary cultures of these cells, whole blood, peripheral mononuclear cells, leukocytes, or immortalized cell lines. Therefore, comparison between studies to identify common key molecular determinants can lead to inconsistent results. Then, compared to studies on more common diseases such as cancer, most of CF transcriptomic studies have very few samples per condition (disease and control), decreasing the statistical power of these datasets when analyzed alone. Finally, these studies rely on various experimental biological models and transcriptomic technologies which rarely lead to consistent results between studies (Clarke et al., 2013), particularly when the analyses are performed at the gene level.

To try and overcome these limitations, we focused on studies considering only samples from human Airway Epithelial Cells (hAEC hereafter), i.e., bronchial, tracheal, or nasal cells. Indeed, functional modifications in these cells are expected to reflect some of the most severe symptoms in the lung. We included studies of cell lines or primary cultures, in order to gather a statistically significant number of samples, because as shown in Table 1, each dataset comprises a very limited number of samples. We also focused on studies on the F508del mutation, for which most data are available. We discarded two studies ((Virella-Lowell et al., 2004) and (Rehman et al., 2021)) that provide transcriptomic data for other mutations, because the corresponding cells could display disparities with respect to F508del cells. We retrieved from the literature all the CF transcriptomic studies with publicly available data that matched these criteria (see Methods section), which led to 10 CF transcriptomic datasets shown in Table 1.

**Table 1.**
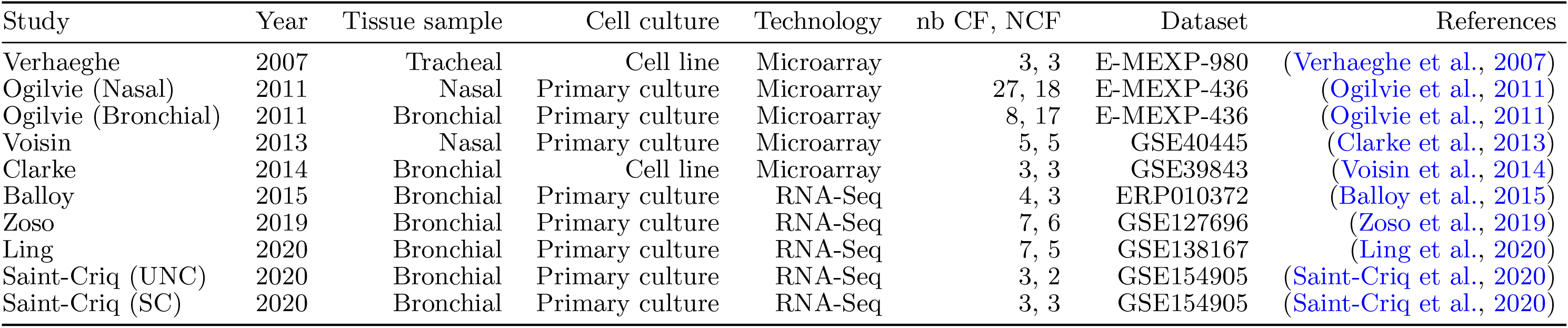
List of the 10 datasets considered in the meta-analysis, indicating the number of CF and NCF samples in each study.

| Study | Year | Tissue sample | Cell culture | Technology | nb CF, NCF | Dataset | References |
| --- | --- | --- | --- | --- | --- | --- | --- |
| Verhaeghe | 2007 | Tracheal | Cell line | Microarray | 3, 3 | E-MEXP-980 | (Verhaeghe et al., 2007) |
| Ogilvie (Nasal) | 2011 | Nasal | Primary culture | Microarray | 27, 18 | E-MEXP-436 | (Ogilvie et al., 2011) |
| Ogilvie (Bronchial) | 2011 | Bronchial | Primary culture | Microarray | 8, 17 | E-MEXP-436 | (Ogilvie et al., 2011) |
| Voisin | 2013 | Nasal | Primary culture | Microarray | 5, 5 | GSE40445 | (Clarke et al., 2013) |
| Clarke | 2014 | Bronchial | Cell line | Microarray | 3, 3 | GSE39843 | (Voisin et al., 2014) |
| Balloy | 2015 | Bronchial | Primary culture | RNA-Seq | 4, 3 | ERP010372 | (Balloy et al., 2015) |
| Zoso | 2019 | Bronchial | Primary culture | RNA-Seq | 7, 6 | GSE127696 | (Zoso et al., 2019) |
| Ling | 2020 | Bronchial | Primary culture | RNA-Seq | 7, 5 | GSE138167 | (Ling et al., 2020) |
| Saint-Criq (UNC) | 2020 | Bronchial | Primary culture | RNA-Seq | 3, 2 | GSE154905 | (Saint-Criq et al., 2020) |
| Saint-Criq (SC) | 2020 | Bronchial | Primary culture | RNA-Seq | 3, 3 | GSE154905 | (Saint-Criq et al., 2020) |

The studies are still heterogeneous in terms of tissue sample (bronchial, tracheal nasal), cell culture type (cell-line or primary culture) and transcriptomic technol- (micro-array or RNA-Seq). However, we kept the 10 studies in order to improve statistical significance, because the numbers of samples per condition are very small in Studies: the median number of samples was 5 for disease (CF) and control (NCF) Conditions.

### 2.3 Meta-analysis of transcriptomic studies at the level of biological pathways

The most straightforward way to analyse transcriptomic data is to identify Differally Expressed Genes (DEGs), and to search for biological pathways enriched in e DEGs. This approach failed in the present meta-analysis, because the number EGs common to at least 3 out of 7 studies was too small to be enriched in any hway, even though many reference pathway databases were considered (the Hall- k gene sets from the the MSigDB Database (Liberzon et al., 2015), the Pathway raction Database (PID) (Schaefer et al., 2009), the KEGG database (Kanehisa l., 2021)). In fact, it has become clear that, in complex diseases, identification of hway dysregulations based on DEGs is not optimal and does not provide robust lts (Wang et al., 2010).

Therefore, the meta-analysis was conducted at the pathway level. Many methods e been proposed to capture pathway dysregulations when they do not appear rly based on enrichment from lists of DEGs (Martignetti et al., 2016; Landais and ot, 2023; Schubert et al., 2018; Vaske et al., 2010). In the present study, we used Gene Set Enrichment Analysis (GSEA) (Subramanian et al., 2005) approach. A was performed separately on each dataset identified as over-activated or undervated signalling pathways in hAEC CF cells, based on the complete expression rix of CF and NCF samples, and taking into account the expression level of all genes belonging to the same pathway. We used pathway definitions provided by the KEGG pathway database (Kanehisa et al., 2021), because this database provides graphical pathway representations that also include phenotypes, which helped the analysis of the CF network, as detailed in Section 2.5. We tested 131 KEGG biological pathways, and Differentially Expressed Pathways (DEPs, hereafter) were identified according to a adjusted p-value lower or equal to 0.25, as detailed in Section 3.4. The number of up- and down-regulated pathways for each dataset is provided in Table 2

**Table 2.** Number of detected genes, CF and NCF samples, tested pathways and dysregulated pathways per study with a corrected p-value *<* 0.25 and a |*log*2*FC*| *>* 1 thresholds at the gene scale.

| Study | nb detected genes | nb CF, NCF | nb up-regulated pathways | nb down-regulated pathways |
| --- | --- | --- | --- | --- |
| Verhaeghe | 22880 | 3, 3 | 11 | 2 |
| Ogilvie (Nasal) | 19880 | 27, 18 | 0 | 0 |
| Ogilvie (Bronchial) | 19880 | 8, 17 | 33 | 0 |
| Voisin | 13144 | 3, 3 | 10 | 0 |
| Clarke | 14118 | 5, 5 | 40 | 2 |
| Balloy | 39430 | 4, 3 | 24 | 2 |
| Zoso | 18846 | 7, 6 | 2 | 0 |
| Ling | 28138 | 7, 5 | 1 | 6 |
| Saint-Criq (UNC) | 39434 | 3, 2 | 11 | 0 |
| Saint-Criq (SC) | 39432 | 3, 3 | 1 | 9 |

The analysis of DEPs showed that 15 of the 134 biological pathways tested were differentially expressed in at least 3 studies. However, a closer analysis highlighted discrepancies between studies. As shown in the heatmap presenting the GSEA Normalised Enrichment Score (NES) (Figure 2), for these 15 common DEPs, the 10 datasets can be gathered into 2 subgroups: subgroup 1 comprising 7 datasets in which common DEPs tend to be up-regulated, while they tend to be down-regulated in subgroup 2 comprising the 3 other datasets. This appeared somewhat puzzling. Our hypothesis is that datasets belonging to subgroups 1 or 2 arise from studies in which the differentiation media used for the primary cultures did not favor the same cell type, and therefore, should not be analyzed together.

**Fig. 2.**
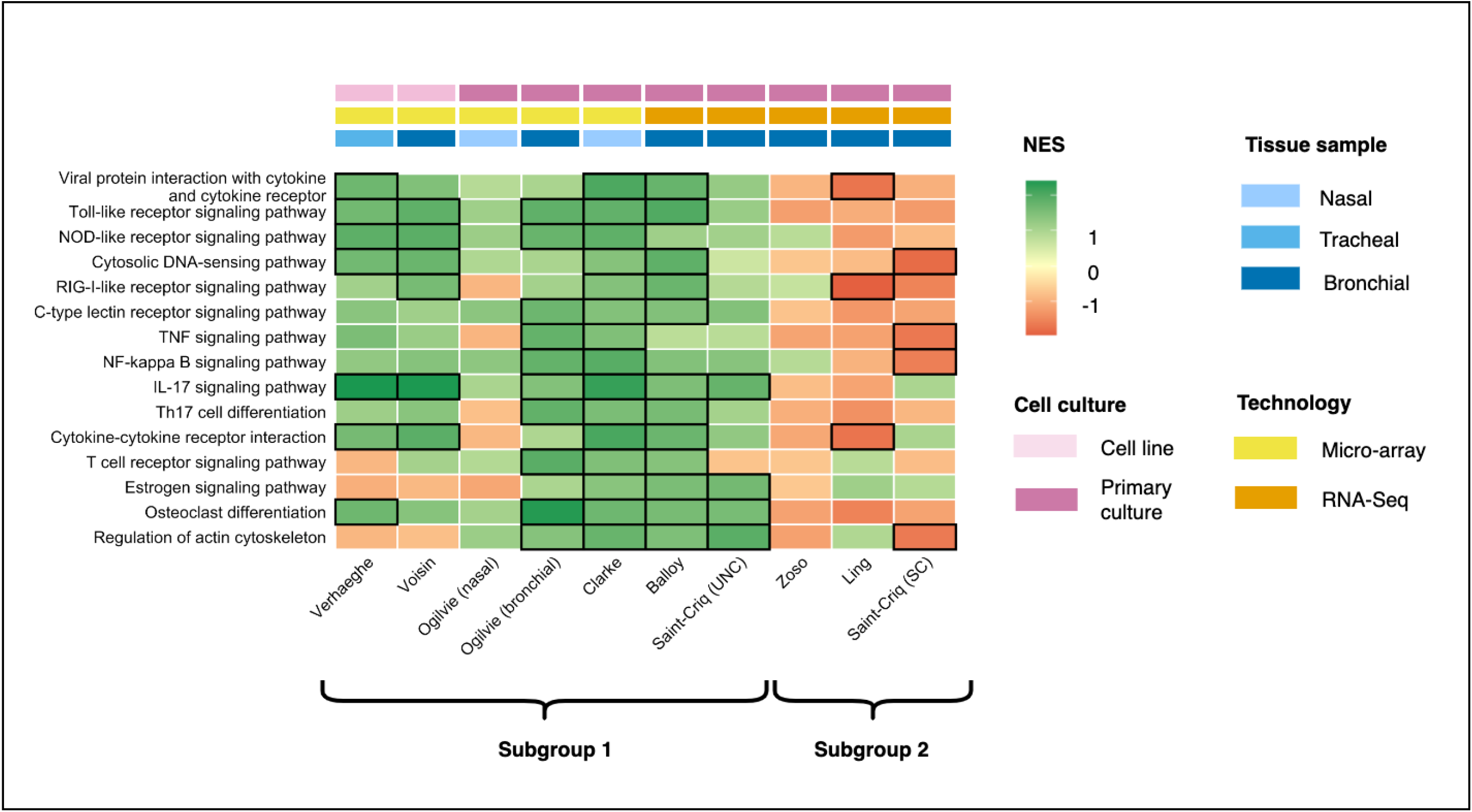
Heatmap of the GSEA Normalized Enrichment Scores (NES) of the biological pathways differentially expressed in at least 3 studies. The datasets can be clustered in two subgroups based on their NES: Subgroup 1 and Subgroup 2, respectively in agreement and in contradiction with CF physio-pathology. Black boxes around the tiles represent the pathways significantly differentially expressed in the corresponding dataset.

This was confirmed by the Saint-Criq (UNC) and Saint-Criq (SC) datasets (see Table 1), belonging respectively to subgroups 1 and 2, where it was shown that the UNC and SC differentiation media (two common differentiation media used on CF and non-CF epithelia) significantly impact cell lineage in primary cultures of CF hAEC, and consequently, the resulting transcriptomic profiles (Saint-Criq et al., 2020). In this study, it was shown that the UNC medium promoted differentiation into club and goblet cells, while the SC medium favored the growth of ionocytes and ciliated cells. Consistent with this result, the Ling transcriptomic dataset, which belongs to subgroup 2, was also obtained from primary cultures of CF and NCF airway epithelia that were differentiated into ciliated pseudo-stratified airway cells (Ling et al., 2020).

Datasets from subgroup 2 appeared in contradiction with the main CF phenotypes. In particular, the *TNF-α signalling pathway* or *NF-κB signalling pathway* are down- regulated in this subgroup, although the over-activation of these pathways is a well- known feature of CF disease. Therefore, we only considered the 7 datasets belonging to subgroup 1 for further analysis.

In this subgroup, the transcriptomic analysis appears to be highly consistent, since among the 15 DEPs common to at least 3 studies, 5 are up-regulated in CF vs NCF samples in 4 studies (*NOD-like receptor signalling pathway, Cytosolic DNA- sensing pathway, Cytokine-cytokine receptor signalling pathway*, and *Regulation of actin cytoskeleton*), 2 are up-regulated in CF vs NCF samples in 5 studies (*Osteoclast differentiation* and *Toll-like receptor signalling pathway*), and the *IL-17 signalling pathway* is up-regulated in CF vs NCF in 6 studies.

Overall, the 15 DEPs common to at least 3 studies are in agreement with various known aspects of CF disease, which confirms that our analysis did capture relevant information about CF. In particular, besides the TNF-*α* and NF-*κ*B signalling path- ways well known to be up-regulated in CF, the IL-17 pathway contributes to CF lung disease (Hsu et al., 2016), the differentiation of osteoclast is perturbed in CF (Dumortier et al., 2021), the Toll-like receptor signalling pathway modulates function, inflammation and infection of lung in CF (Kosamo et al., 2020; Curutiu et al., 2018; Fleurot et al., 2022), and CFTR plays a role in cell junction and actin cytoskeleton organization (Pankonien et al., 2022).

### 2.4 Building the CF network

The 15 individual DEPs of the KEGG database provide interesting information about what is dysregulated in CF, but a lot of these pathways are partially redundant and show a high overlap of genes and interactions, indicating that they are highly intertwined. A dysregulation in one of these pathways will have a consequence in another pathway. To study the connection between them, we propose to merge them into a single network called the CF network.

The DEPs were extracted with the OmniPathR package (Tü rei et al., 2016) and curated, as described in Section 3.5. The rules that were used to build and clean this network are detailed in Section 3.6. The network, comprising 330 nodes and 529 interactions, is not fully connected: it contains one main component including 317 nodes connected by 515 interactions, and two small additional components that are non connected to the main component, and called unconnected components hereafter (See Figure 3). The overall network can be accessed as a Cytoscape session, in the sysbio-curie/CFnetwork cystoscape github repository for further analysis.

**Fig. 3.**
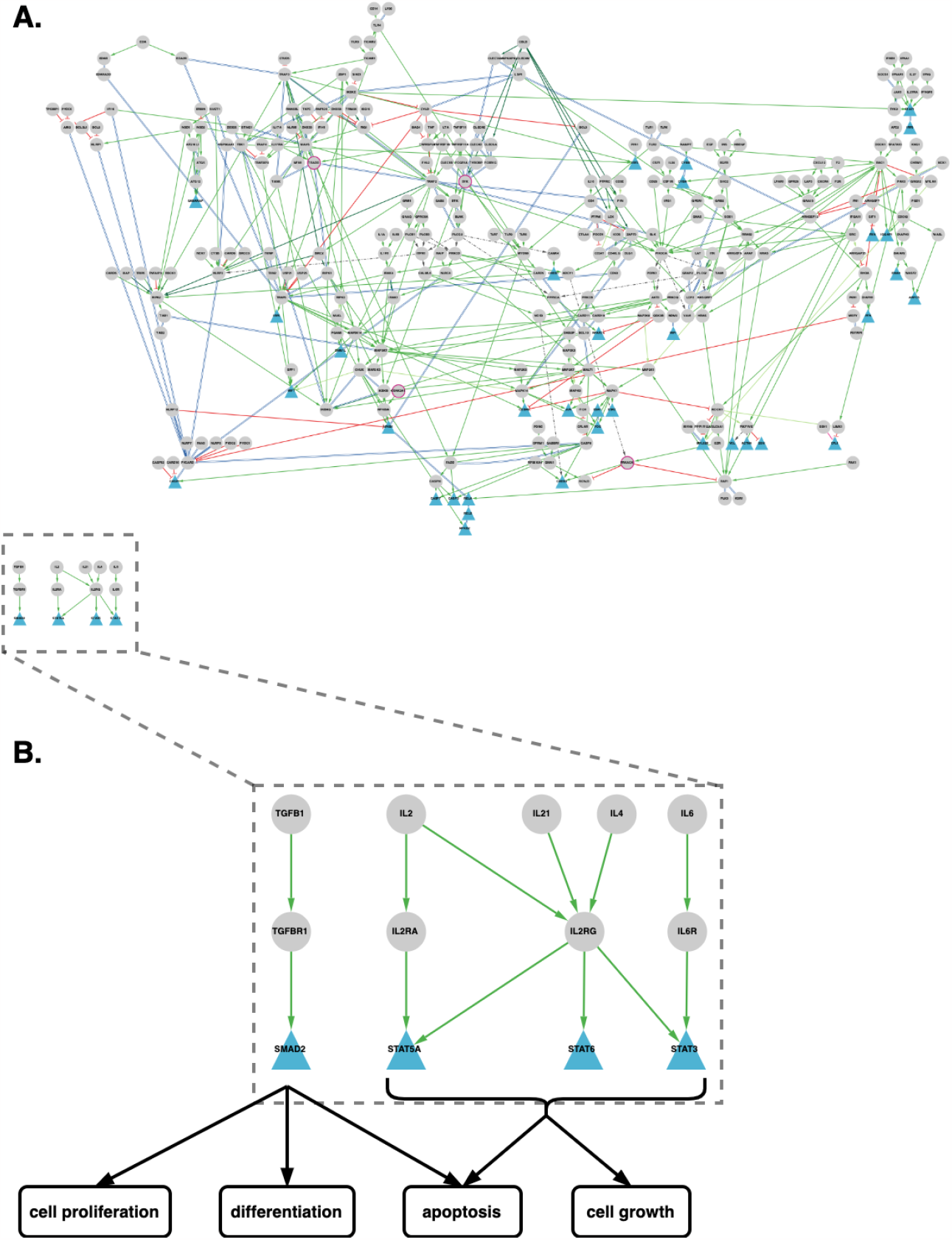
The CF network. (A): The main component comprises 317 nodes connected by 517 interactions and two small unconnected components shown in (B): the two unconnected components correspond to the TGF*β* and the JAK-STAT signalling pathways. The cellular phenotypes triggered by the sink nodes of the two components are surrounded by black contours.

#### Identification of CFTR interactors in the CF network

It is striking to note that CFTR does not belong to any of the 15 DEPs, and there- fore, is not part of the network. In fact, CFTR is present in only 7 biological pathways of the KEGG database (*ABC Transporters, cAMP signalling pathway, AMPK signalling pathway, tight junction, Gastric and acid secretion, pancreatic secretion* and *bile secretion*), but these pathways did not belong to the DEPs.

Therefore, we searched for the presence of CF network proteins in the network of proteins reported to be involved in protein-protein interactions (PPI) with wt-CFTR or F508del-CFTR (Pereira et al., 2021). Indeed, according to the CyFi-MAP, 4 direct interactors of wt-CFTR but not of F508del-CFTR (CSNK2A1, PRKACA, SYK and TRADD) belong to the CF network. Furthermore, 4 additional proteins (EZR, SRC, PLCB1/3) present in the network interact with wt-CFTR (but not with F508del- CFTR) through a single intermediate protein. Figure 4 shows these 8 proteins, their intermediates and their interactions with CFTR. The presence in the CF network of 8 first or second neighbours in the CFTR interactome is an interesting result in favour of our assumption that CFTR interactors may propagate functional dysregulations into the network.

**Fig. 4.**
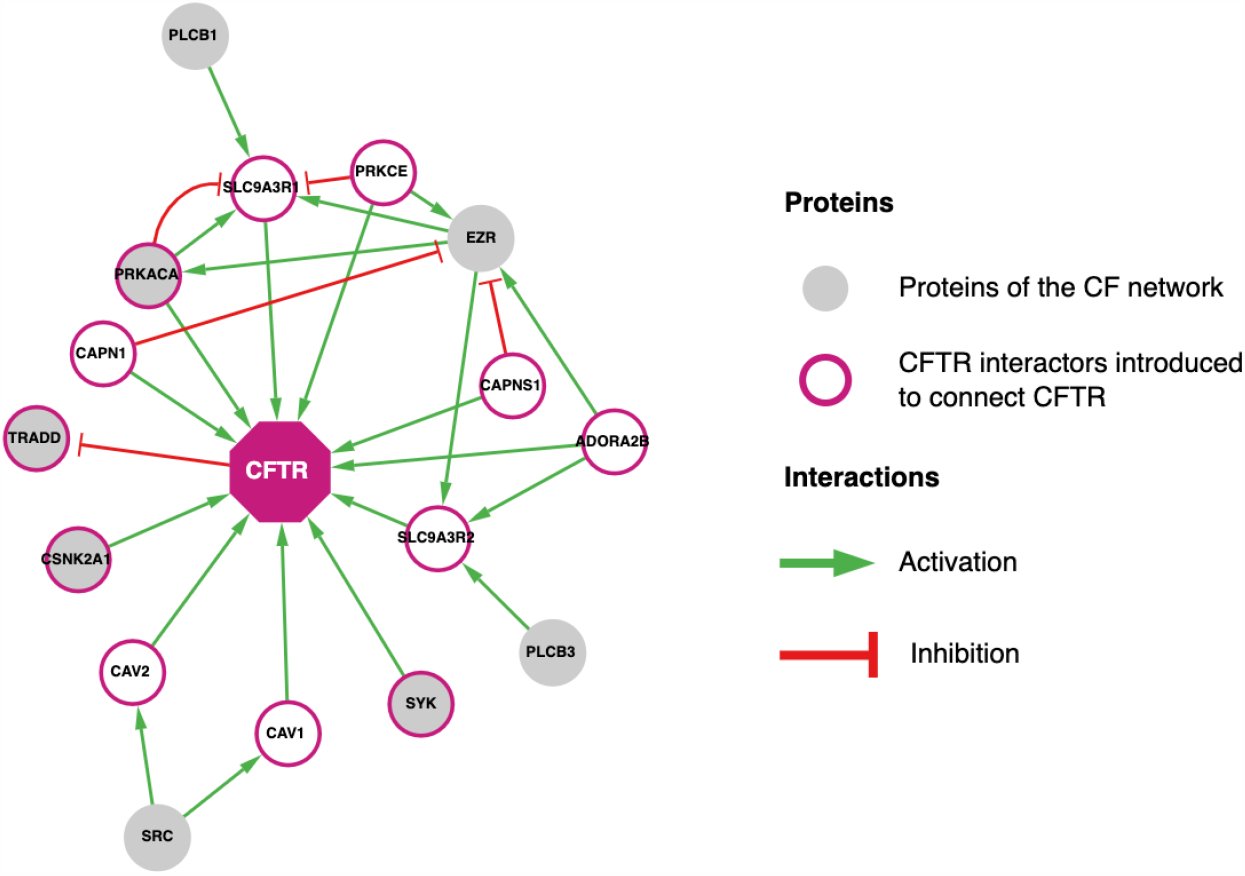
CFTR interactors in the CF network: Known protein-protein interactions involving CFTR interactors in CFTR PPI.

### 2.5 Analysis of the CF network

Extensive interpretation of this large network, which contains rich but complex information, is beyond the scope of the present paper. However, we will investigate how analysis of its topology can help tackle the two questions of interest: how the absence of the CFTR protein at the membrane leads to CF cellular phenotypes, and how therapeutic targets can be suggested from this network.

#### 2.5.1 Topological description of the CF network

The final CF network comprises 330 proteins and 529 interactions. Interestingly, CFTR interactors are present only in the main component, because according to the CyFi- MAP, it would not have been possible to link CFTR to proteins of the two small unconnected components without adding a large number of intermediate nodes. One of the two unconnected components contains 10 proteins and 9 interactions, and corresponds to cascades of the JAK/STAT signalling pathway. The other contains 3 proteins and 3 interactions, and corresponds to a cascade of the Transforming Growth Factor Beta (TGF*β*) signalling pathway. In the present section, we will focus on the main component of the CF network, and the two unconnected components will be discussed in Section 2.5.3.

The topological description of the main component will be organized around three types of remarkable nodes: (1) the source nodes, i.e., CFTR first or second neighbours that were used to connect CFTR to the network, as described in Section 2.4; (2) the sink nodes, i.e., the nodes from which no edge leaves in the network, and whose activation finally triggers their associated phenotypes (for example, transcription factors are typical sink nodes); (3) the hubs, i.e. the nodes with high betweenness centrality, through which the flow of information that passes is high. Figure 5 illustrates where these remarkable nodes stand within the network’s topology.

**Fig. 5.**
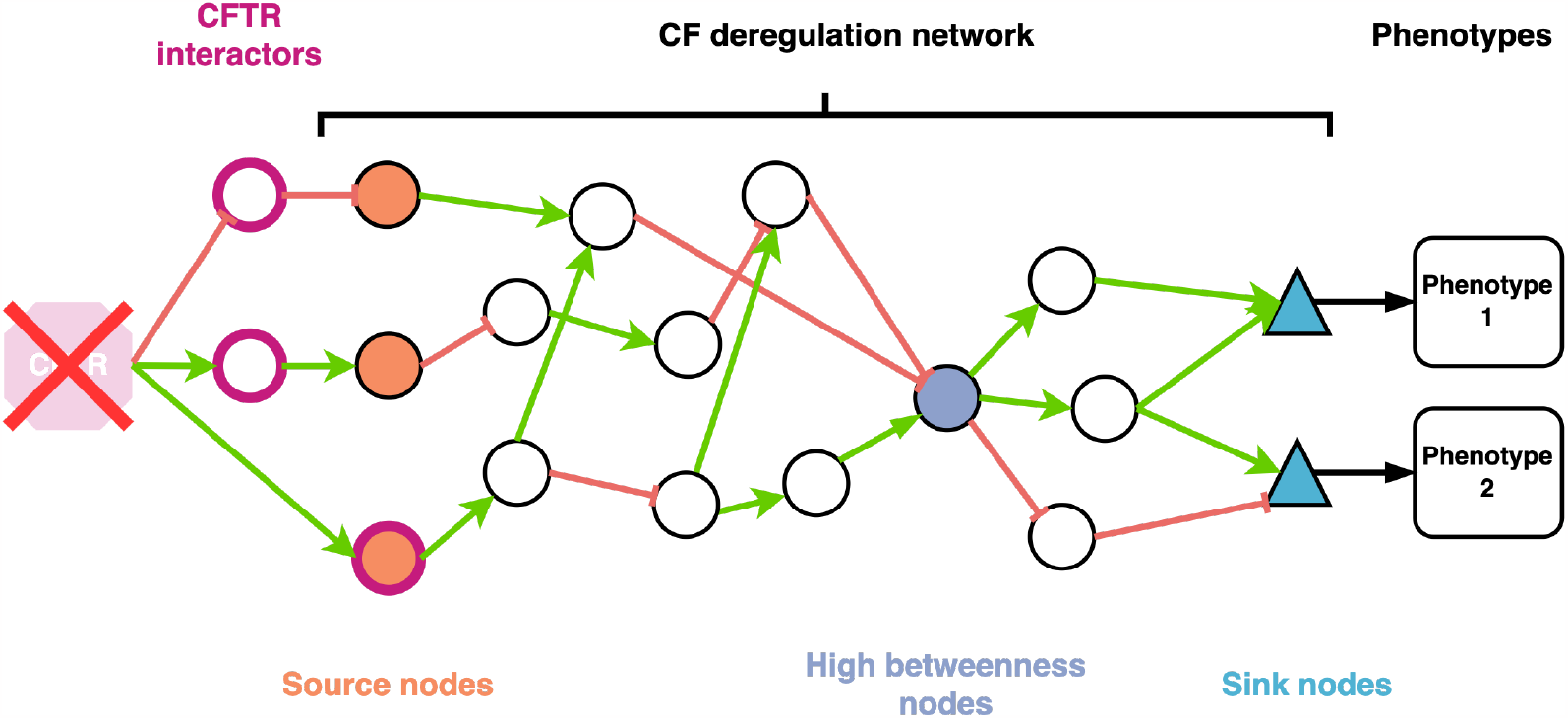
Illustration of propagation of dysregulation and remarkable nodes in the CF network: the source nodes (orange disks) are CFTR interactors or connected to CFTR interactors via a single intermediate protein (magenta circles). Nodes with high betweenness centrality (purple disks) are proteins through which much information flows within the network. Sink nodes (blue triangles) modulate their corresponding phenotypes.

##### Source nodes and initiation of dysregulations

According to the CyFi-MAP, 8 first or second neighbours of wt-CFTR interactors whose interactions are lost with F508del-CFTR are present in the CF network: **CSNK2A1, EZR, PLCB1, PLCB3**,**PRKACA, SRC, SYK** and **TRADD**. In the absence of CFTR, these 8 proteins can be viewed as source nodes that may initiate dysregulations that subsequently propagate within the network and finally reach the sink nodes (see Figure 4).

Perturbations of some of these source nodes in CF cells, or their role in CF cellular phenotypes, are sustained by various studies:

- **CSNK2A1**, also known as CK2 (casein kinase 2), is strongly overactivated in CF vs wild-type cells (Venerando et al., 2011).
- Cellular levels of **TRADD** are controlled by its lysosomal degradation in a wt- CFTR-dependant manner, and this regulation is lost with F508del-CFTR and G551D-CFTR (Wang et al., 2016).
- **SRC** was shown to be overexpressed and overactivated in CF cells (Massip Copiz and Santa Coloma, 2016).
- **PLCB3** is a known CF modifier gene, for which the loss of function S845L variant is associated with a mild progression of the pulmonary disease and a reduction of *Pseudomonas aeroginosa*-induced IL8 release. This indicates that PLCB3 plays a role in the inflammation phenotype in CF (Rimessi et al., 2018).
- The active form of ezrin (**EZR**) is mainly located in the apical region of wild type airway epithelial cells, while in their CF counterparts, it is diffusely expressed in its inactive state in the cytosol (Favia et al., 2010; Wu and Eickelberg, 2019).
- The **SYK** and **PRKACA** kinases play key roles with respect to CFTR, since the former negatively regulates the amount of CFTR at the membrane through phosphorylation at Y512 (Mendes et al., 2011), while the latter is a well-known regulator of the CFTR chloride channel conductance (Egan et al., 1992), but their implication as propagators of dysregulations has not been investigated yet.

##### Sink nodes and CF phenotypes

There are 35 sink nodes in the main component of the CF network that are reached from each of the 8 source nodes. The full list of sink nodes and their associated phenotypes are given in the Supplementary file. Among them, we can cite:

**NFKB1, NFKB2, RELA** and **RELB** are part of the NF-*κ*B complex, a transcription factor that can be activated by various stimuli such as cytokines, oxidant radicals, bacterial or viral products. It controls the expression of pro-inflammatory genes, and is related to various phenotypes including inflammation and cell survival/proliferation.

**FOS** and **JUN** are two sub-units of the AP-1 transcription factor activated by the MAPK signalling pathways, and are associated with inflammation and proliferation phenotypes.

**CASP3** and **CASP7** caspases are the effectors of apoptosis.

**CASP1** is a caspase known to be the effector of pyroptosis, a highly proinflammatory cell death mechanism.

10 sink nodes belong to the regulation of actin cytoskeleton pathway, including **ACTN4, ARPC5, PFN, MYL12B** and **VCL**. These nodes are associated to various phenotypes related to cytoskeleton, including focal adhesion, adherens junction, and actin polymerisation.

**IRF1, IRF3, IRF5**, and **IRF7**, that are members of the IRF family of transcription factors involved in the innate immune response phenotype, and controlling expression of Type-1 interferons upon viral infection.

Importantly, the phenotypes associated to these sink nodes have already been described in the CF context. In particular:

1. The NF*κ*B and AP-1 transcription factors are complexes of sink nodes that mediate inflammation, the most studied phenotype of CF disease. In addition to the well-known activation of NF*κ*B in CF, AP-1 is one of the downstream transcription factors of the MAPK pathway that was shown to be activated in CF (Bérubé et al., 2010; Wellmerling et al., 2022), as shown in Figure 6.
2. Controversial results were reported about apoptosis in CF epithelial cells. Some studies showed defective susceptibility of CF cells to pro-apoptotic stimuli (Cannon et al., 2003; Gottlieb and Dosanjh, 1996), while others observed increased apoptosis (Chen et al., 2018; Voisin et al., 2014; Yalçcin et al., 2009; Rottner et al., 2007). All agree that apoptosis is dysregulated in CF.
3. The dysregulation of actin cytoskeleton in CF is well documented, with a disorganized actin cytoskeleton, absence of actin stress fibres (Favia et al., 2010; Lasalvia et al., 2016; Burat et al., 2022), and disrupted tight junctions (De Lisle, 2014; Castellani et al., 2012).
4. Finally, most of these phenotypes are related to the innate immune response and various works indicate a dysfunction in the innate immune response of CF patients (Kosamo et al., 2020; Gillan et al., 2023; Dugger et al., 2020).

**Fig. 6.**
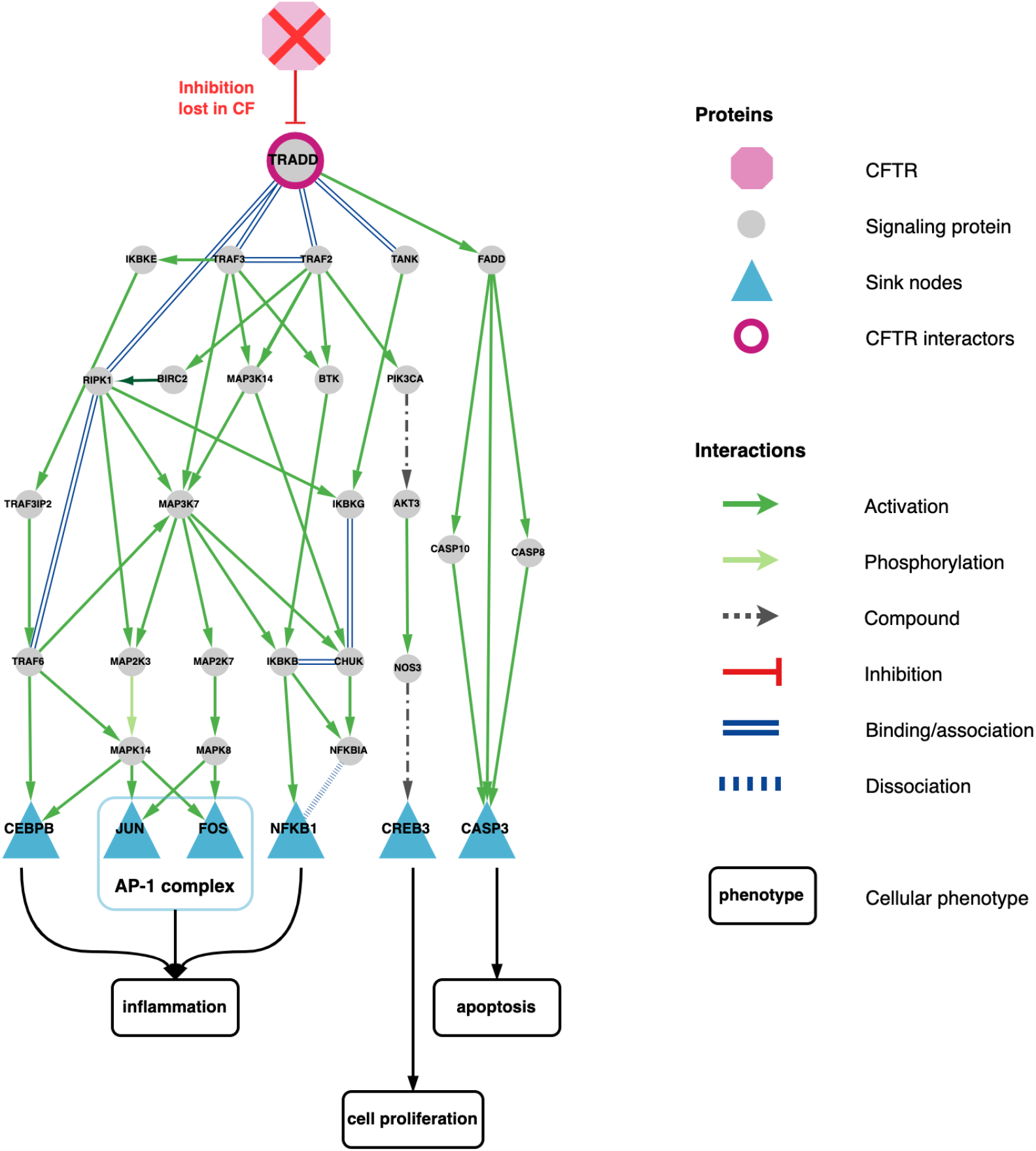
Extract from the CF network showing the TRADD protein connected to the TNF-*α* signalling pathway, and to 5 other sink nodes, including FOS and JUN which form the AP-1 transcription factor, downstream of the MAPK cascade. The cellular phenotypes triggered by the sink nodes are surrounded by black contours. Note that TRADD is connected to the 35 sink nodes, but only part of the nodes downstream of TRADD in the network are represented.

##### Betweenness centrality and flow of information

In a network, the betweeness centrality (BC) of a node is the number of shortest paths that pass through that node. This measure is a way of detecting the amount of influence a node has over the flow of information in a network. Nodes with high BC, referred to as hubs, may provide interesting therapeutic targets, because their inhibition may efficiently reduce the propagation of information within the network (Durón et al., 2019). Therefore, we calculated the BC for all nodes of the CF network, as detailed in the Methods section. All nodes were then ranked according to this measure, and Figure 7A displays the histogram of the BC score. Interestingly, most proteins have a BC score below 3000, and only a very limited number of proteins have a BC score above 6000 (ARHGEF12, IKBKE, LSP1, PIK3KC1, PYCARD, RAC1, TRAF2, TRAF3, TRAF6). The list of the top 30 proteins is provided in the Supplementary File. Among them, PI3KCA could be an interesting therapeutic target candidate and is discussed in the next section.

**Fig. 7.**
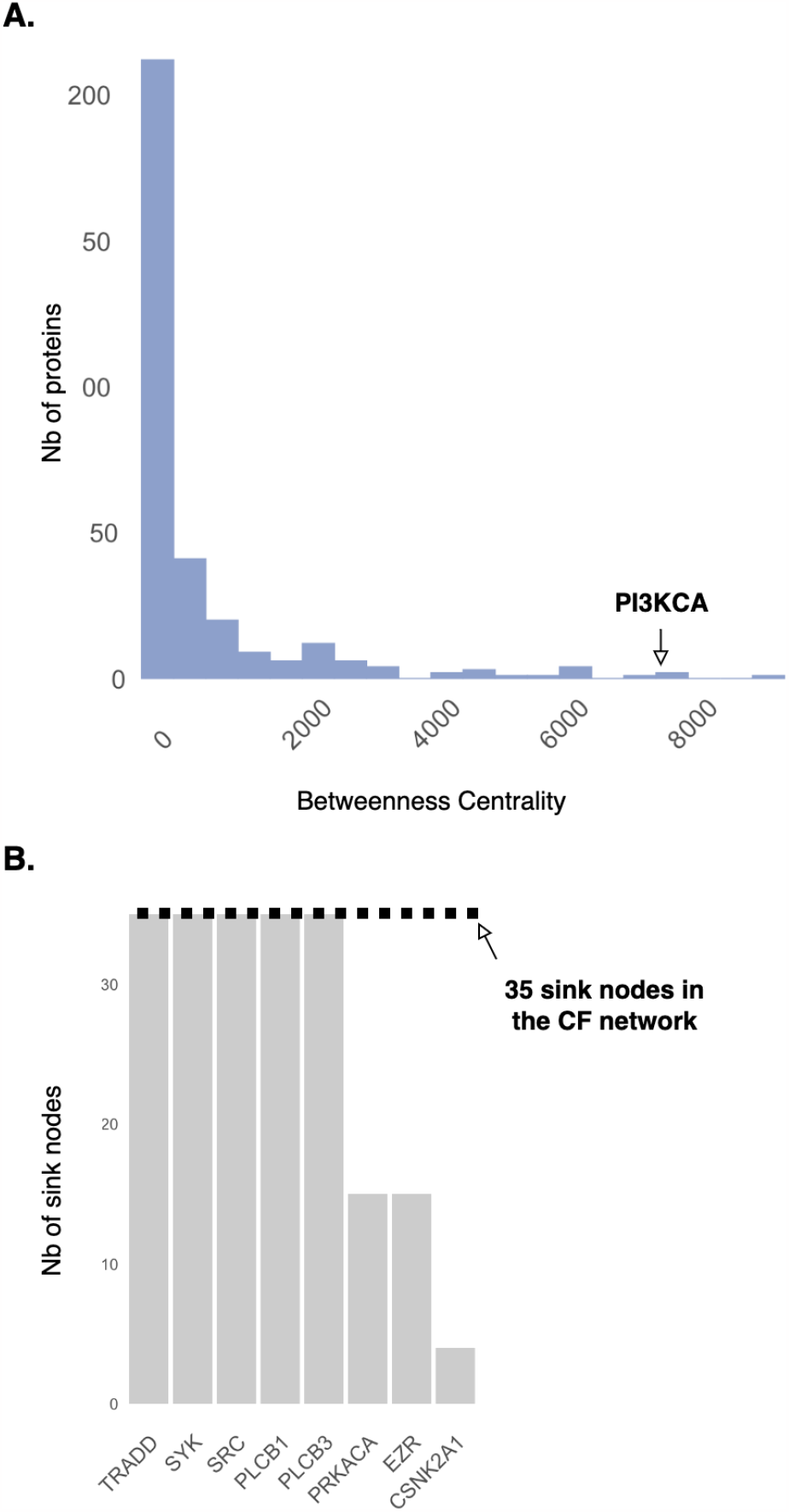
(A) Histogram of the betweenness centrality measures for all nodes in the CF signalling network; (B) Number of sink nodes to which each of the 8 source nodes are connected.

#### 2.5.2 Biological insights from the topological analysis

A simple path analysis of the network shows that several source nodes may contribute collectively to the emergence of the CF phenotypes, which illustrates the complexity of the disease. Indeed, while the source nodes PRKACA, EZR and CSNK2A1 are upstream of a limited number of sink nodes, PLCB1/3, SRC, SYK and TRADD are upstream of the 35 sink nodes, i.e., there exists a path from each of these 6 source nodes to each of the 35 sink nodes (Figure 7B).

For example, TRADD is known to be up-regulated in CF (Ferenc Karpati, 2000) and to participate in the uncontrolled inflammation (See Figure 6). Interaction between wt-CFTR and TRADD enhances the degradation of TRADD, which controls the activity of this pathway, as demonstrated by Wang and colleagues (Wang et al., 2016). This direct interaction is lost with F508del-CFTR, which may contribute to the dysregulation of TNF-*α* and NF-*κ*B signalling pathways in CF. However, up- regulation of TRADD could also contribute to the inflammation phenotype through another route, by inducing over-activation of the MAPK pathway, and in particular of AP-1, one of its output transcription factors. In addition, as shown in Figure 8, our network suggests that other source nodes than TRADD could also initiate dysregulation of the inflammation phenotype because they are also connected to the NF-*κ*B sink node. Among these sources, we can cite: (1) SYK, which would be consistent with its role in inflammation processes shown in other diseases (Riccaboni et al., 2010; Wong et al., 2004); (2) PLCB1/3, which are consistent with previous studies reporting PLCB3 as a key modulator of IL8 expression in CF bronchial epithelial cells (Bezzerri et al., 2011); (3) CSNK2A1, whose hyperactivity could contribute to activation of NF-*κ*B by enhancing the phosphorylation and degradation of IKBKA.

**Fig. 8.**
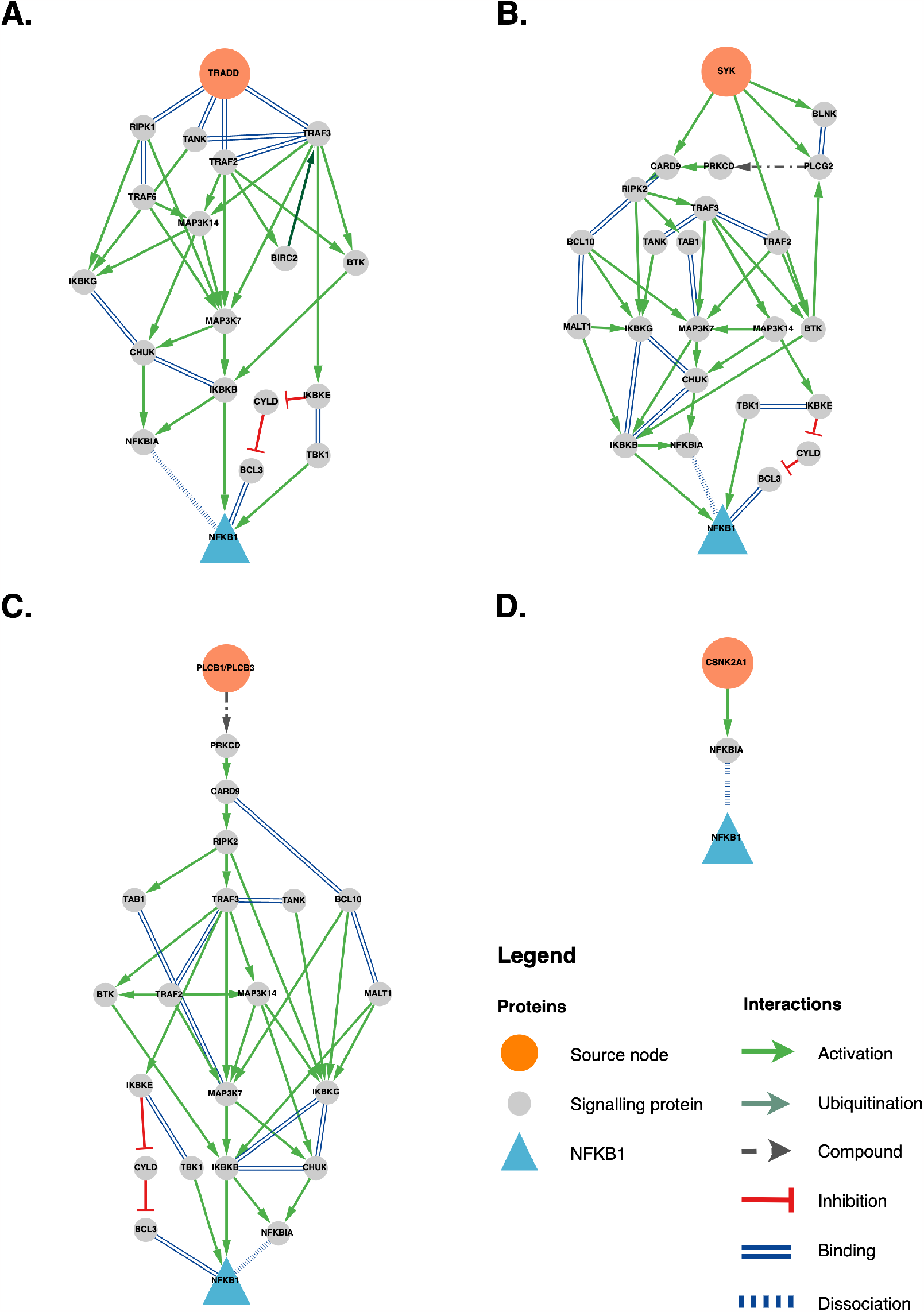
Subnetworks of the CF network illustrating the connections between the source nodes TRADD, SYK, PLCB1/3, and CSNK2A1 and the sink node NFKB1.

Overall, the number of source nodes and routes that may contribute to inflammation in CF illustrates the challenge posed by its modulation, in order to reduce the related clinical symptoms. Various anti-inflammatory drugs have been recently evaluated in clinical trials (Bell et al., 2020), but none of them target the source nodes of the present study. Our hypothesis is that these source nodes could be interesting candidate targets in CF. In particular, SYK has emerged as a potential target for the treatment of numerous diseases. Many inhibitors are known for this kinase, which would allow to evaluate their potential anti-inflammatory effect in CF cells. These inhibitors include one marketed drug (Fostamatinib), but other inhibitors are currently under investigation in clinical trials for a range of indications (Cooper et al., 2023). Interestingly, since SYK is connected to the 35 sink nodes, its inhibition may also contribute to the modulation of other CF phenotypes than inflammation. In particular, it could modulate CF phenotypes associated to the 35 sink nodes and mentioned in Section 2.5.1, such as dysregulations in apoptosis, cytoskeleton or innate immune response. Similarly, our network suggests that PLCB3 could be an interesting target for inflammation in CF. This is consistent with the fact that PLCB3 silencing in CF bronchial epithelial cells exposed to *Pseudomonas aeroginosa*, reduces the expression of IL-8 chemokine (Bezzerri et al., 2011). The U73122 PLC inhibitor could be an interesting pharmacological tool to further evaluate this strategy. As in the case of SYK, PLCB3 is connected to the 35 sink nodes, which means that its inhibition may also improve other CF cellular phenotypes. Consistent with this idea, it was shown that treatment with a SRC inhibitor, another of the 6 source nodes upstream of the 35 sink nodes, decreased the inflammatory changes and improved cytoskeletal defects in F508del human cholangiocytes (Fiorotto et al., 2018).

Besides source nodes, candidate therapeutic targets can be searched among hubs in the network, i.e. among the best ranked proteins according to the BC score. Besides this score, additional arguments can be invoked to highlight the best candidates. In particular, the fact that a protein is known in the literature to play a role in the disease, and that pharmacological modulators (or even better, marketed drugs) are available to allow experimental validation, are important criteria. In line with these ideas, PI3KCA appears as an interesting candidate target. Indeed, several inhibitors are known for this kinase, including the marketed drug Alpelisib, which would allow experimental tests in CF models. It has been suggested as a candidate target in CF based on its role in many signalling pathways implicated in CF lung pathogenesis (Natarajan, 2020). The fact that PI3KCA belongs to best ranked proteins with respect to the BC score (See Figure 7A) offers a quantitative argument in favor of this idea. In addition, PI3KCA is connected through the network to the 35 sink nodes, which means that its inhibition may modulate inflammation, but also other CF cellular phenotypes related to the sink nodes.

#### 2.5.3 Analysis of the unconnected components in the CF network

As mentioned in Section 2.5.1, Figure 3 shows that the CF network comprises two small unconnected components that are part of the TGF*β* and JAK/STAT signalling pathways. Contrary to source nodes of the main component, dysregulation of the source nodes of these unconnected components (namely the 4 interleukins IL2, IL21, IL4 and IL6 for one component, and TGF*β* for the other) cannot be explained by the absence of CFTR in a direct manner, because they are not linked to CFTR within a single network. However, activation of a sink node of the main component may modulate the expression of a source node in an unconnected component, affecting the activity of this unconnected component. For example, activation of the AP-1 transcription factor (a sink node of the main component) due to activation of the MAPK pathway in the main component, regulates the expression of TGF*β*. This example shows how dysregulations in one pathway may have consequences in other pathways of the CF network, even if they are not connected, again illustrating to the complexity of the disease. We propose that phenotypes arising from the two unconnected components could be defined as secondary phenotypes, as opposite to primary phenotypes arising from dysregulations of the main component (discussed in Section 2.5.1).

The JAK-STAT component mediates various cellular processes, including cell growth and apoptosis, but the role of these cascades has not been widely studied in CF. The TGF*β* component leads to the activation of SMAD2, a transcriptional modulator that regulates multiple cellular phenotypes, including cell proliferation, apoptosis, and differentiation. High levels of TGF*β* have been associated with the severity of lung disease (Dorfman et al., 2008; Sagwal et al., 2020), and this protein was proposed as a therapeutic target for CF (Kramer and Clancy, 2018). Our study suggests that therapeutic targets should be chosen among proteins closer to CFTR in the network, in particular among the source nodes of the main component (as discussed above), because they may more successfully limit the global propagation of molecular dysregulations within the overall network.

## 3 Methods

### 3.1 Datasets selection

Based on the search engines of the National Center for Biotechnology Information (NCBI) and the European Nucleotide Archive (ENA), we selected 10 datasets from 8 studies published between 2007 and 2021. The selection criteria to include CF transcriptomic datasets were the following: (1) they should correspond to human Airway Epithelial Cells (hAEC); (2) the cells should be homozygous for the most common mutation F508del; (3) the transcriptomic data should be publicly available. Therefore, studies including samples heterozygous for the F508del mutation ((Virella-Lowell et al., 2004) and (Rehman et al., 2021)), studies with no data available (Zabner et al., 2005) and (Wright et al., 2006)) were not included. In addition, studies with less than two samples were excluded ((Bampi et al., 2020) and (Veltman et al., 2021)), as the subsequent statistical analyses require several samples per condition. The list of selected transcriptomic studies is provided in Table 1.

### 3.2 Biological pathways databases

We initially considered a total of 380 gene sets corresponding to 380 biological path- ways: 50 Hallmark gene sets from the the Molecular Signatures Database (MSigDB) (Liberzon et al., 2015), 196 from the Pathway Interaction Database (PID) (Schaefer et al., 2009) and 134 from the KEGG database, restricted to the *Genetic Information Processing, Environmental Information Processing* ; *Cellular Processes* and *Organismal systems* subdivision. However, most of the analyses were performed using only KEGG database. Indeed, in the Hallmark and the PID databases, gene sets are defined as gene signatures rather than as biological pathways. Thus, the genes are not necessarily connected to each other through functional interactions. Conversely, gene sets retrieved from the KEGG database correspond to biological pathways defined as genes corresponding to proteins that participate in oriented molecular cascades. They are available in the form of maps on the KEGG website. In addition, the structure of the KEGG database allows to build a network that provides mechanistic interpretation. Therefore, gene set enrichment algorithms required to build the signalling network was performed based on the KEGG database. All interactions and nodes from each biological pathway of the KEGG database were retrieved thanks to the OmnipathR R package (Tü rei et al., 2016).

### 3.3 Preprocessing of RNA-Seq data

Limma was originally developed for differential expression analysis of microarray data, which values are assumed to be normally distributed, and the variance independent of the mean. This is not the case for log2-counts per million (log-CPM) values in RNA-Seq data: expression distributions may vary across samples and methods modelling counts assume a quadratic mean-variance relationship. Therefore, for the RNA-Seq data, 3 steps of pre-processing are necessary before applying the statistical tests (Law et al., 2018): (1) low expressed genes are filtered (i.e. genes with less than 10 read counts in at least one sample in the condition with the minimum sample size); (2) normalisation using the method of trimmed mean of M-values (TMM) is performed (Robinson and Oshlack, 2010); (3) raw counts are converted to log-CPM and the mean-variance relationship is estimated with the *voom* method.

### 3.4 Identification of Differentially Expressed Pathways (DEPs)

For each of the 10 transcriptomic datasets, identification of DEPs was performed using the *fgseaSimple* function of the Bioconductor package *fgsea* (Korotkevich et al., 2021), for fast preranked Gene Set Enrichment Analsyis (GSEA) (Subramanian et al., 2005). The *fgseaSimple* method takes two inputs: a gene-level signed statistics and a defined list of genes known as *gene set*. The method ranks the genes in descending order based on the chosen statistics, and then computes the Enrichment Score (ES) for the gene set. The ES reflects how often members of that gene set occur at the top (e.g., upregulated) or the bottom (e.g., downregulated) of the ranked gene list. To account for differences in gene sets size, a normalisation step is performed to obtain the Normalised Enrichment Score (NES). Besides, random gene sets are generated and their NES computed. These NES are then used to create a null distribution from which the significance of the NES of the tested gene set is estimated. In our study, we used the t-statistics from the differential expression analysis comparing gene expression levels of CF sample to NCF samples as the control condition. In order to compare all the studies together, all the microarray and RNA-Seq datasets were processed using the same pipeline, involving the limma (Ritchie et al., 2015) and edgeR (Robinson et al., 2010) packages. After removing technical outlier samples and retrieving gene symbols using the biomaRt package (Durinck et al., 2009), differential expression analysis at the gene level was performed by fitting a linear model using weighted least squares for each gene.

Gene sets with size larger than 500 were excluded for statistical testing. The p-values of the gene sets were adjusted for multiple testing error with Benjamini- Hochberg (BH) procedure. Differentially Expressed Pathways (DEP)s were considered with a corrected p-value lower or equal to 0.25. If the NES is positive, the DEP is categorized as up-regulated, and if it is negative, the DEP is categorized as down-regulated.

#### 3.5 Up-dating Omnipath DEPs pathways

The CF network was built from DEPs among pathways in the KEGG database, as extracted with the OmniPathR package. We observed a few inconsistencies between the corresponding list of genes and interactions downloaded with OmnipathR R package, and those in the ’up to date’ pathways maps, as they are displayed on KEGG website. Therefore, we updated the OmnipathR version of the KEGG pathways by adding (or removing) a few nodes or interactions, in order to map the OmnipathR pathways with their corresponding pathways in KEGG. For each modification, bibliographic references were manually checked into other databases stored in Omnipath, in particular in the high confident databases SignorDB (Lo Surdo et al., 2022), and the Human Reference Interactome (Drew et al., 2021). In addition, in a few pathways, some interactions are labelled as ”indirect” in KEGG database. They involve part of signalling cascades belonging to other biological pathways, and they are not detailed in the considered pathway. For example, part of the PI3K-AKT pathway belongs to the Toll-like receptor signalling pathway but is not detailed in this pathway (See KEGG map for Toll-like receptor signalling pathway). In such cases, in order to build the network based on complete cascades involving only direct interactions, we added the missing nodes and interactions.

All the pathways modifications and the corresponding codes used to perform these modifications are available in the following github repository: sysbio-curie/CFnetwork.

### 3.6 Network building and pruning

In the KEGG database, most of the 15 common DEPs display the same overall topology: some cell-surface receptor proteins activate one or more intra-cellular signalling cascades that in turn activate downstream transcription factors, thus triggering corresponding phenotypes. For example, the NF-*κ*B pathway leads to the ”inflammation” or ”cell survival” phenotypes. However, 2 of the common DEPs, *Cytokine-cytokine receptor interaction* and *Viral protein interaction with cytokine and cytokine receptor*, are pathways that do not consist in such functional cascades. The *Cytokine-cytokine receptor interaction* pathway consists in a list of interactions between extra-cellular signal molecules and cell-surface receptors (see KEGG database to visualise this pathway’s topology). These interactions are also part of larger biological pathways that comprise their corresponding downstream cascades. This means that KEGG pathways are partially redundant (i.e. small pathways are part of larger pathways), which is also found in all commonly used pathway databases. In the case of the *Cytokine-cytokine receptor interaction* pathway, this DEP is dysregulated in the meta-analysis because some of the interactions between extracellular molecules and cell surface receptors are dysregulated, but not necessarily all of them. For example, interactions between TNF-*α* and its receptors, or IL17 and its receptors are dysregulated, but this information is also present in the DEPs containing the complete corresponding cascades, i.e. the *TNF-α signalling pathway* and the *IL-17 signalling pathway*. The same type of analysis also holds for the *Viral protein interaction with cytokine and cytokine receptor* DEP. Overall, from these 2 DEPs, we only retained the cell-surface receptors that are sources of downstream dysregulated cascades in our network. Overall, 25 cell surface receptors without downstream dysregulations in our CF transcriptomic data were removed from the network.

Finally, we also removed from the pathways all the interactions corresponding to genes targeted by transcription factors, downstream of the pathways’ cascades, because these target genes do not define the pathways themselves.

Network building and pruning were performed using the R packages tidyr v.1.2.1, and dplyr v.1.0.10. Transcription factors were identified using the R packages Dorothea v.1.4.2 and hgnc v.0.1.2, which give access to the Dorothea (Garcia-Alonso et al., 2019) and HUGO collections (Seal et al., 2023), respectively.

### 3.7 Betweenness centrality score

The betweenness centrality (BC) score of node *n* is defined by

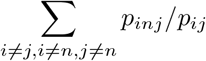

where *p*_*ij*_ is the total number of shortest paths between nodes *i* and *j* while *p*_*inj*_ is the number of those shortest paths which pass though vertex *n*.

BC scores were computed using the *betweenness* function of the R package igraph v.1.3.4 (Csardi and Nepusz, 2005). This pacakge was also used for the other network topology analyses.

### 3.8 Network Visualization and Figure Generation

The networks, generated as dataframes in R, were imported into Cytoscape v.3.9.0 (Shannon et al., 2003) for visualization. We designed a custom style for nodes and edges, which is available in the Cytoscape session and also saved as an independent XML file, available in the sysbio-curie/CFnetwork cystoscape github repository. The hierarchical layout was used to emphasize the information flow from the source nodes to the sink nodes.

Barplots were generated using the R package ggplot2 v.3.3.6, and didactic figures were created using the open-source platform diagrams.net.

## 4 Discussion

Using a pathway-based meta-analysis of publicly available transcriptomic data, we built the CF network that provides a more global understanding of the molecular dysregulations in CF than the view of a CFTR-related channelopathy disease. Indeed, an important outcome of this work was to integrate data analyses to network reconstruction, while proposing a strategy to relate CFTR to proteins of the network, based on CFTR interactome. The CF network comprises a restricted number of source nodes that connect the absence of CFTR to the downstream sink nodes triggering CF cellular phenotypes. Another important contribution was to propose candidate therapeutic targets, based on the topological analysis of this network (namely, SYK, PI3KCA and PLCB1/3). The network provides a comprehensive view of how pathway interactions contribute to a given disease phenotype. It reveals unintuitive effects of targeting candidate proteins because of the complex interactions of the biological pathways in the network. Overall, the CF network can be seen as a tool to formulate hypotheses and interpret experimental observations.

Although several transcriptomic datasets were gathered, the total number of samples globally included remains modest (57 CF and 46 control samples). Additional data may refine the list of dysregulated pathways, and help to improve the proposed CF network.

To cope with the low number of samples per study, we opted for a meta-analysis combining various CF transcriptomic datasets, which highlighted that distinct differentiation media used for the primary cultures may favor different cell types, leading to inconsistent transcriptomic profiles and potential erroneous interpretations. This may explain why previous transcriptomic comparative studies reported incoherent signs of gene dysregulation (up- versus down-) between different datasets for many genes (Clarke et al., 2013). We observed the same phenomenon at the pathway level for datasets belonging to subgroup 1 or 2 (see Section 2.3). Clustering studies based on the heatmap of common DEPs appears to be a good tool to select consistent data in future meta-analysis.

Other types of dysregulations such as aberrant phosphorylations are not detectable in transcriptomic data. Including information from other types of omic data such as proteomic, phosphoproteomic, metabolomic,or volatilomic may help to refine the CF network. In particular, in the past three years, CF airway epithelial single-cell RNAseq (sc-RNAseq) datasets have been reported (Carraro et al., 2021; Thurman et al., 2022). Such data allow the study of dysregulations at the cell type level, and could facilitate building of the CF networks for specific epithelial cell types. Furthermore, CFTR is expressed in cell types beyond airway epithelial cells. Thus, refining this network within the context of these cell types could enhance our understanding of the role of CFTR in these specific cells such as macrophages, where CFTR seems to have non-channel functions (Duan et al., 2021).

Prior knowledge gathered in the KEGG pathway database was used to identify and connect DEPs, but the proposed methodology can be followed using other pathways databases. Pathway names and definitions vary between databases, and therefore, the resulting network may slightly depend on the reference database that was used. Nevertheless, it would comprise globally the same interactions and proteins. Similarly, CFTR interactors present in the network were identified according to PPI information in the CyFi-MAP. If new CFTR interactors are identified, this information may help improve the content of the network, highlighting new source nodes or routes for the propagation of dysregulations. In particular, missing interactions, because they are not present in pathway databases, or have not been discovered yet, may explain the presence of unconnected components. If they exist, their discovery in the future may allow to link the two unconnected small components to the main component of the network. However, the proposed notion of targeting proteins as upstream as possible in the network, or among key hubs of the network, are still an interesting concept in order to prioritize candidate therapeutic targets.

An important issue of the present paper was to explore the link between absence of the CFTR protein, and more global pathway dysregulations that lead to CF cellular phenotypes. However, the precise definition of a diseased cellular phenotype is not clearly defined yet, and we used key words provided in the KEGG database or in the Gene Cards database (Stelzer et al., 2016). The present work proposes an answer this question in the context of systems biology studies. Associating phenotypes to the activity of outputs of the signalling cascades, referred to here as sink nodes, could be a first step towards the definition of the disease read-outs. This is of particular interest for *in vitro* evaluation of drug candidates, because we expect that drugs active in CF would reduce the activity of these sink nodes.

The methodological approach proposed in our study was settled based on transcriptomic data from hAEC cells homozyguous for F508del, because publicly available data are more abundant for this most frequent mutation. Therefore, our CF network characterizes the disease caused by this mutation. It would be interesting to study to which extent the CF network would differ for other mutations. A recent paper indicates that DEGs in human bronchial epithelial cell lines bearing mutations from different classes share about 30% DEGs, while 70% of the DEGs are class specific (Santos et al., 2023). It would be interesting to study if this still holds at the level of biological pathways, as they are defined in the present work, and to study whether the resulting network is strongly modified, or not. The methodology proposed in the present paper and based on network topology could still be applied in order to search for new, and potentially class-specific, therapeutic targets.

The candidate therapeutic targets proposed based on our CF network could also be tested on CF cellular models for other mutations, because these targets may belong to biological pathways that are also dysregulated with other mutations. If this was the case, it would help to extend the therapeutic arsenal available for CF patients who are not eligible for CFTR modulators.

In the same line, it is now clear that CF patients bearing the same mutation may present diseases of different severity. Although many factors can modulate disease severity, including environmental factors, it would be interesting to explore the contribution of patients molecular profiles. In particular, building a ”personalized” CF network based on patients’ transcriptomic profiles would be an interesting tool to answer this question.

Beyond CF, reduced amounts of functional CFTR have also been observed in other diseases like chronic obstructive pulmonary disease (COPD) (Saint-Criq and Gray, 2017; Simões et al., 2021), cigarette smoke (Valdivieso et al., 2018), or cancer (Duan et al., 2021; Wang et al., 2022). The network could provide a basis to explore the consequences of reduced CFTR activity in these diseases.

## 5 Conclusion

We presented building of **the CF network**, a signalling network gathering the molecular dysregulations caused by the absence of CFTR. We adopted a data-driven systems biology approach to retrieve CF dysregulated signalling pathways. These pathways were merged to build a signalling network, recapitulating the dysregulated cascades that flow from the source nodes (proteins connected to CFTR) to the sink nodes (proteins that trigger CF cellular phenotypes). Five of the source nodes are upstream of all the sink nodes in the CF network: PLCB1/3, TRADD, SRC,and SYK. These proteins may collectively initiate the emergence of CF phenotypes (together with the other 3 source nodes EZR, CSNK2A1, and PRKCA), illustrating the complexity of the disease. The topological analysis of the network also highlighted nodes with a high degree of betweenness centrality, which are other important players in the propagation of the dysregulations, including PI3KCA. Among these key source nodes and nodes with high degree centrality, SYK, SRC, PLCB1/3 and PIK3CA appeared as interesting candidate therapeutic targets. Interestingly, specific inhibitors are known for these proteins, and even marketed drugs in the case of SYK and PI3KCA. They stand out as potential therapeutic candidates for drug repositioning, potentially allowing the modulation of various CF phenotypes. Finally, an important contribution of the present work is that the adopted global methodology of the CFTR context, although perfectible, did provide interesting results for CF, and can be used as a common framework for other monogenic diseases.

**Additional File 1**. CFnetwork additional file 1.pdf, a PDF document with extended results concerning the sink nodes and their corresponding cellular phenotypes:

**Table S1**: The 35 sink nodes of the CF network and their corresponding cellular phenotypes.

**Table S2**: The top 30 proteins in the CF network according to their betweenness centrality scores.

## List of abbreviations

BC: Betweenness Centrality
CF: Cystic Fibrosis
CFTR: Cystic Fibrosis Transmembrane Conductance Regulator
DEG: Differentially Expressed Gene
DEP: Differentially Expressed Pathway
GSEA: Gene Set Enrichment Analysis
hAEC: human Airway Epithelial Cells
NCF: Non-cystic Fibrosis
NES: Normalised Enrichment Score
PPI: Protein-Protein Interaction

## Declarations

## Ethics approval and consent to participate

Not applicable

## Consent for publication

Not applicable

## Availability of data and materials

The codes and datasets supporting the conclusions of this article are available in the folowing github repository: sysbio-curie/CFnetwork.

The Cytoscape session of the CF network, the TSV files of the nodes and the edges of the CF network and the XML file of the custom style of the Cytoscape session required to reproduce the Cytoscape session are available in the following github repository: sysbio-curie/CFnetwork cystoscape.

## Author’s contribution

I.S. and V.S. initiated the project and obtained funding for the project. M.N., L.M., L.C. and V.S. contributed to the conception and design of the research. M.N. and L.M. designed the transcriptomic analysis. M.N. conducted the transcriptomic analyses, built the signalling network and performed network analyses under the supervision of L.C. and V.S.. M.C., M.K.A. and I.S. contributed to the interpretation of the data. M.N., L.C. and V.S. wrote the manuscript. L.M. I.S. and M.K.A edited the manuscript. All authors read and approved the final manuscript.

## Acknowledgments

The authors would like to than Benoît Chevalier and Alexandre Hinzpeter for fruitful discussions on CFTR interactome, and Marco Ruscone for fruitful discussions on biological databases.

