## Additional File for "From CFTR to a CF signalling network: a systems biology approach to study Cystic Fibrosis"

### Table S1. The 35 sink nodes of the CF network and their corresponding cellular phenotypes.

| HGNC | Cellular phenotypes |
|---|---|
| CASP1 | pyroptosis, cell death, inflammation |
| CASP3 | apoptosis |
| CASP7 | apoptosis |
| CYBA | ROS/oxidative stess |
| CYBB | ROS/oxidative stess |
| DNM1L | Necroptosis/Cell Death |
| GABARAP | Autophagy |
| ACTN4 | Regulation of actin polymerisation |
| ARPC5 | Regulation of actin polymerisation |
| CFL1 | Regulation of actin polymerisation |
| ENAH | Regulation of actin polymerisation |
| GSN | Regulation of actin polymerisation |
| IQGAP1 | Regulation of actin polymerisation |
| MYL12B | Regulation of actin polymerisation |
| PFN | Regulation of actin polymerisation |
| PXN | Regulation of actin polymerisation |
| VCL | Regulation of actin polymerisation |
| CEBPB | inflammation |
| CREB1 | cell cycle, apoptosis, inflammation |
| CREB3 | proliferation, migration, differentiation, inflammation |
| ESR1 | Regulation of cell cycle, apoptosis, cell adhesion |
| ESR2 | Regulation of cell cycle, apoptosis, cell adhesion |
| FOS | innflammation, proliferation |
| IRF1 | innate immune response |
| IRF3 | innate immune response |
| IRF5 | innate immune response |
| IRF7 | innate immune response |
| IRF9 | innate immune response |
| JUN | inflammation, proliferation |
| NFATC1 | cellular differentiation, immune response |
| NFKB1 | inflammation, cell survival/proliferation |
| NFKB2 | inflammation, cell survival/proliferation |
| RELA | inflammation, cell survival/proliferation |
| RELB | inflammation, cell survival/proliferation |
| STAT1 | innate immune response |

Cellular phenotypes were retrieved from the KEGG database or from the Gene Cards database when no phenotype was associated with the sink node in any of the KEGG pathways.

### Table S2. The top 30 proteins in the CF network according to their betweenness centrality score

| HGNC | BC score |
|---|---|
| TRAF2 | 9541.17 |
| LSP1 | 7987.59 |
| PYCARD | 7845.23 |
| PIK3CA | 7710.44 |
| IKBKE | 6734.15 |
| TRAF3 | 6691.04 |
| TRAF6 | 6365.37 |
| ARHGEF12 | 6268.20 |
| RAC1 | 6018.87 |
| MAVS | 5527.33 |
| STING1 | 5000.21 |
| IFI16 | 4890.92 |
| PAK3 | 4813.73 |
| TBK1 | 4456.73 |
| MAP3K7 | 4345.00 |
| SYK | 3622.03 |
| VAV1 | 3347.09 |
| ZAP70 | 3309.76 |
| PLCG2 | 3295.18 |
| TRAF3IP2 | 3086.74 |
| LCP2 | 2974.65 |
| CLEC7A | 2944.23 |
| RHOA | 2830.65 |
| NLRP3 | 2814.41 |
| AKT3 | 2784.62 |
| CASP8 | 2748.34 |
| IKBKG | 2723.52 |
| HSP90AA1 | 2705.86 |
| MAPK1 | 2657.41 |
| CYLD | 2628.59 |
